# Deciphering the transcriptomic landscape of tumor-infiltrating CD8 lymphocytes in B16 melanoma tumors with single-cell RNA-Seq

**DOI:** 10.1101/800847

**Authors:** Santiago J. Carmona, Imran Siddiqui, Mariia Bilous, Werner Held, David Gfeller

## Abstract

Recent studies have proposed that tumor-specific tumor-infiltrating CD8^+^ T lymphocytes (CD8 TIL) can be classified into two main groups: “exhausted” TILs, characterized by high expression of the inhibitory receptors PD-1 and TIM-3 and lack of transcription factor 1 (Tcf1); and “memory-like” TILs, with self-renewal capacity and co-expressing Tcf1 and PD-1. However, a comprehensive definition of the heterogeneity existing within both tumor-specific and total CD8 TILs has yet to be clearly established.

To investigate this heterogeneity at the transcriptomic level, we performed paired single-cell RNA and TCR sequencing of CD8 T cells infiltrating B16 murine melanoma tumors, including cells of known tumor specificity. Unsupervised clustering and gene signature analysis revealed four distinct CD8 TIL states - exhausted, memory-like, naïve and effector memory-like (EM-like) - and predicted novel markers, including Ly6C for the EM-like cells, that were validated by flow cytometry. Tumor-specific PMEL T cells were predominantly found within the exhausted and memory-like states but also within the EM-like state. Further, TCR repertoire sequencing revealed a large clonal expansion of exhausted, memory-like and EM-like cells with partial clonal relatedness between them. Finally, meta-analyses of public bulk and single-cell RNA-seq data suggested that anti-PD-1 treatment induces expansion of EM-like cells.

Our reference map of the transcriptomic landscape of murine CD8 TILs will help interpreting future bulk and single-cell transcriptomic studies and may guide the analysis of CD8 TIL subpopulations in response to therapeutic interventions.

## INTRODUCTION

Chronic antigen stimulation and inflammation lead to CD8 T cell differentiation and function that differ from those generated during acute infections. Chronically stimulated CD8 T cells acquire an “exhausted” state characterized by a progressive loss of cytolytic activity, reduced cytokine production and proliferative capacity, upregulation of multiple co-inhibitory receptors, such as PD-1, CTLA4, LAG3, TIGIT and TIM3, and a unique epigenetic state (Mclane, Abdel-Hakeem and Wherry, 2019). Although initially considered hypofunctional effector T cells, several observations suggest that exhausted T cells are heterogeneous and have crucial roles in limiting viral infection or tumor growth while avoiding damage to normal tissues (Speiser *et al.*, 2014). Yet, antigen-specific T cells in tumors often lack effector function and fail to control tumor growth and therefore they are also referred to as “dysfunctional” (Li *et al.*, 2019; Philip and Schietinger, 2019). Notwithstanding, immune-checkpoint blockade (ICB) therapies using anti-PD-1 can result in a proliferative response of CD8 tumor-infiltrating lymphocytes (TILs) and improve effector functions (Tumeh *et al.*, 2014).

Understanding how anti-PD-1 therapy affects distinct CD8 TIL subsets is a major challenge in cancer immunotherapy. Recently, a novel intratumoral tumor-specific CD8 T cell subpopulation was discovered among murine TILs that mediates cellular expansion in response to immune checkpoint blockade (Miller *et al.*, 2019; Siddiqui *et al.*, 2019). These cells have been isolated and characterized using different combinations of surface markers and reporter genes (e.g. PD-1^+^ Tcf7:GFP^+^ and PD-1^+^ TIM3^-^ SLAMF6^+^) and were named “memory-like” (Siddiqui *et al.*, 2019) or “progenitor exhausted” (Miller *et al.*, 2019). Memory-like CD8 TILs cells have the capacity to expand, self-renew and give rise to terminally differentiated cells (PD-1^+^ TIM3^+^ Tcf1^-^, GZMB^+^; termed “exhausted”) in the context of chronic antigenic stimulation, similarly to progenitor CD8 T cell subsets previously described in chronic infection (He *et al.*, 2016; Im *et al.*, 2016; Utzschneider *et al.*, 2016; Wu *et al.*, 2016; Kallies, Zehn and Utzschneider, 2019). However, there are still many unknowns regarding how T cells committed to the exhaustion lineage develop from pre-exhausted T cell states and which of these states are present in the tumor. Therefore, a comprehensive definition of the heterogeneity existing within both tumor-specific and total CD8 TILs has yet to be clearly established.

Here we aimed at defining a reference transcriptomic map and determining clonal relatedness of CD8 TILs, including cells of known tumor-specificity, in the common B16 murine melanoma model. To this aim, we have sequenced the transcriptome including full-length T cell receptor genes of >3500 single-cell CD8 TILs from seven tumor-bearing mice including wild-type C57BL/6 and PMEL transgenic mice. Furthermore, to study how the CD8 TIL landscape is modulated by immunotherapy, we performed a meta-analysis of published bulk and single-cell RNA-seq (scRNA-seq) data. Our study provides new insights into the heterogeneity of CD8 TILs, including novel markers to define T cell subpopulations, as well as freely accessible bioinformatics tools to guide the analysis of scRNA-seq data sets.

## RESULTS

### Single-cell RNA-seq of CD8 TILs reveals the presence of exhausted, memory-like, naïve and effector memory-like T cells

To obtain an unbiased view of the transcriptomic landscape of tumor-infiltrating CD8 T cells from B16 melanoma tumor-bearing mice, we performed single-cell transcriptomic profiling paired with VDJ locus sequencing of CD8 TILs. We individually analyzed 4 wild-type (WT) C57BL/6 mice and 3 PMEL transgenic mice, whose CD8 T cells express a transgenic TCR specific for the B16 tumor-associated antigen gp100/PMEL (Figure 1 A). After data processing and quality control, >3500 CD8 T cells from the 7 tumors were kept for downstream analyses (See Methods). Unsupervised clustering on the high-dimensional space revealed the presence of four robust CD8 TIL clusters with distinct transcriptomic profiles. Cluster 1 (C1) was defined by high expression of inhibitory receptors *Pdcd1, Havcr2, Ctla4, Tigit* and *Lag3*, exhaustion-related transcription factors such as *Batf* and *Tox* (Alfei *et al.*, 2019; Khan *et al.*, 2019; Scott *et al.*, 2019; Yao *et al.*, 2019) and high expression of cytotoxic molecules (e.g. *Gzmb, Prf1, Fasl*), compatible with an exhausted state (Figure 1 B,C). Cluster 2 (C2), in proximity to C1, was defined by the co-expression of inhibitory receptors (expressing *Pdcd1, Tigit* and *Lag3* but not *Havcr2* or *Entpd1*) and memory related genes (e.g. *Tcf7, Lef1, Sell*), and low levels of cytotoxicity genes (Figure 1 B,C), compatible with the recently described memory-like subset (Miller *et al.*, 2019; Siddiqui *et al.*, 2019). Cluster 3 (C3) was defined by high expression of markers of naïve/memory (e.g. *Tcf7, Sell, Ccr7, Lef1, Il7r*) and no expression of cytotoxicity genes or inhibitory receptors, compatible with a naïve or central memory state (Figure 1 B, C). Based on the lack of *Cd44* expression in this cluster (Figure 1 C) we provisionally refer to it as naïve cells cluster. Finally, cells in cluster 4 (C4) expressed high levels of memory-related genes (e.g. *Tcf7, Lef1, Il7r*) together with cytotoxicity genes (e.g. *Gzmk, Gzmb*). Compared to the memory-like C2, C4 was characterized by low expression of the inhibitory receptors *Pdcd1, Tigit, Lag3* and the exhaustion-related transcription factors *Tox* and *Batf* (Figure 1 C). Expression of granzymes and lack of *Sell* and *Ccr7* expression suggested an effector memory rather than a central memory phenotype (Figure 1 C). Hence, this cluster was referred to as Effector Memory-like (“EM-like”) state (Figure 1 B,C). Note that, while T cells with effector memory phenotype have been previously observed among TILs (Zhang *et al.*, 2018; Haas *et al.*, 2019), EM-like cells have never been characterized at the transcriptomic level on murine tumors, and a gene signature for this population remains to be defined. Differentially expressed genes in each cluster are shown in Supplemental Table 1. Gene expression differences between these four clusters could be further summarized by the expression of three gene sets associated with 1) cytotoxicity (*Gzmb, Prf1* and *Fasl*), 2) inhibition/exhaustion (*Pdcd1, Havcr2, Tigit, Lag3, Ctla4*) and 3) “stemness” (*Tcf7, Sell, Il7r, Lef1*). While the naïve cluster (C3) presented the highest level of stemness and the lowest inhibition and cytotoxicity, the exhausted cluster (C1) presented the lowest levels of stemness with the highest levels of inhibition and cytotoxicity (Figure 1 D). The memory-like (C2) cluster displayed higher levels of “stemness” compared to exhausted, together with lower levels of inhibition and very low levels of cytotoxicity, in line with previous observations (Miller *et al.*, 2019; Siddiqui *et al.*, 2019). Instead, the EM-like (C4) cluster displayed intermediate levels of cytotoxicity and “stemness”, with low levels of inhibition (Figure 1 D).

**Figure 1.**
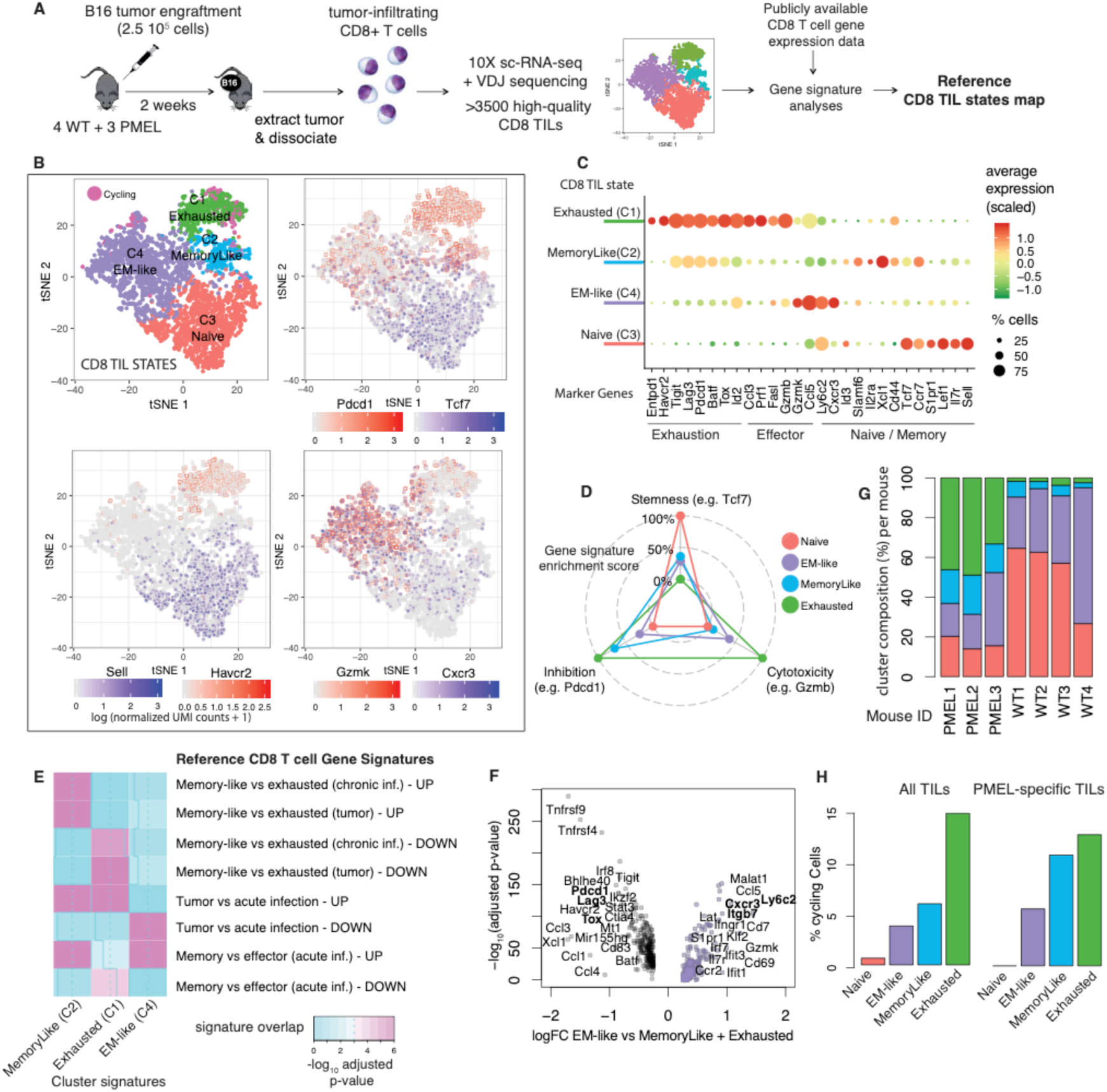
Defining CD8 TIL states. **A** Overview of experimental design. **B** tSNE plots indicating global transcriptomic similarities of CD8 TILs, and colored by unsupervised clustering (upper-left panel, clusters C1 to C4) or by expression of specific marker genes (other panels). Cycling cells are marked in magenta. **C** Dotplot indicating average expression of a panel of marker genes (x-axis, associated with naïve/memory, exhaustion and effector T cell phenotypes) for the four T cell clusters (y-axis). Color scale indicates scaled and centered log-normalized UMI counts. **D** Projection of T cell states onto three phenotypic scores axes (stemness, inhibition/exhaustion and cytotoxicity, see Methods). Phenotypic scores are relative to each other, varying from 0% (inner circle) for the lowest score, to 100% (outermost circle) for the highest score. **E** Gene signature enrichment analysis against reference CD8 T cell subtypes signatures observed in cancer and chronic infection. UP and DOWN refer to the sets of up- or down-regulated genes associated to each comparison. Color scale indicate statistical significance of signature overlap (FDR corrected p-values, Fisher’s exact test). Details about the reference signatures in Methods. **F** Volcano plot showing top differentially expressed genes between EM-like vs exhausted and memory-like states. **G** Relative T cell cluster composition for each mouse. **H** Percentage of cycling cells in each cluster, as defined by high expression levels of cell-cycle genes in all TILs (left) or PMEL-specific TILs (expressing PMEL TCR, right). See methods for details.

We next evaluated to what extent these CD8 TIL states relate to previously described CD8 T cell subsets found in the context of cancer or infection. An initial gene signature enrichment analysis confirmed that C3 matched the transcriptomic state of splenic naïve T cells (Sarkar *et al.*, 2008), while the other three clusters up-regulated genes associated with differentiated CD8 T cells (Supplemental Figure 1, see Methods). Next, we focused on the three differentiated states to evaluate signature enrichment against specific CD8 T cell subtypes. We found a consistent mapping of the memory-like (C2) cluster with the tumor-resident PD-1^+^ Tcf1^+^ “memory-like” subset (Siddiqui *et al.*, 2019) (Figure 1 E), whereas the exhausted cluster (C1) was mapped to the “exhausted” PD-1^+^ Tcf1^-^ subset described in the same study. Further, these two clusters also matched the memory-like and exhausted subsets, respectively, found in chronic infection (Utzschneider *et al.*, 2016) (Figure 1 E).

The EM-like cluster did not match the exhausted nor the memory-like signatures, and instead showed specific enrichment for the signature of pathogen-specific CD8 T cells found upon acute infection (“Tumor vs acute infection – DOWN”, row 6 in Figure 1 E) (Schietinger *et al.*, 2016). Further, among pathogen-specific CD8 T cells found upon acute infection, the EM-like cluster was specifically enriched in the signature of memory (day 60 post LCMV Armstrong infection, “Memory vs effector (acute inf.) – UP”) rather than effector (KLRG1^+^ day 4.5 post LCMV Armstrong infection) CD8 T cells (“Memory vs effector (acute inf.) – DOWN”) (rows 7 and 8 in Figure 1 E). Hence, signature enrichment analysis confirmed an effector memory (EM-like) phenotype for this population. Since EM-like cells have not been previously characterized in B16 tumors, we analyzed this cluster in more detail. Differential gene expression analysis of EM-like vs exhausted and memory-like cells revealed potential novel markers for this population (Figure 1 F), including *Ly6c2* that encodes a surface molecule that has been previously associated to memory CD8 T cells (Walunas *et al.*, 1995; Pihlgren, 1996; Cerwenka *et al.*, 1998) and *Cxcr3*, a chemokine receptor that guides the recruitment of T cells into inflamed peripheral tissue (Groom and Luster, 2011).

Next, we analyzed how CD8 TIL states were distributed among WT and PMEL mice, i.e. according to antigen-specificity. We found that T cell from both types of mice were present in all four states (Figure 1 G and Supplemental Figure 2), although a clear distribution bias was observed. TILs from PMEL mice were enriched in exhausted (33-49% of PMEL vs 2-4% of WT) and memory-like (14-20% of PMEL vs 3-8% of WT) states. TILs from WT mice were enriched in EM-like (17-37% of PMEL vs 26-69% of WT) and naïve (14-20% of PMEL vs 27-65% of WT) T cells. Thus, while tumor-specific cells are enriched in the exhausted and memory-like states, total polyclonal CD8 TILs are enriched in the EM-like and naïve states. Interestingly, our analysis revealed the presence of tumor-reactive (PMEL) cells in the EM-like state.

Cycling cells (i.e. with high expression of cell cycle-related genes) were detected within exhausted (16%), memory-like (6%) and EM-like (4%) states (cells in magenta in Figure 1 B, and Figure 1 H left panel), as opposed to cells from the naïve cluster that did not cycle (<1%). When considering PMEL-specific T cells (effectively expressing PMEL TCR receptor, see Methods), a similar distribution was observed (Figure 1 H, right panel). This indicates that in addition to exhausted and memory-like, EM-like cells, including tumor-specific EM-like cells, replicate in the tumor.

The robustness of the four identified transcriptomic states was confirmed by unsupervised clustering of an independent publicly available scRNA-seq dataset of CD8 TILs from B16 melanoma tumors (Singer *et al.*, 2016), where a consistent cluster correspondence was verified between datasets (Supplemental Figure 3). Furthermore, re-analysis of a recently published scRNA-seq dataset of tumor-specific CD8 TILs in B16-OVA tumors (Miller *et al.*, 2019) revealed the presence of EM-like cells (12% among OVA Tetramer^+^ CD8 TILs) in addition to memory-like (11%) and exhausted (76%) cells (Supplemental Figure 4), further supporting that part of the EM-like subset contains tumor-specific cells.

Overall, we were able to define the landscape of transcriptomic states of endogenous CD8 TILs in B16 melanoma tumors and recapitulate the naïve, memory-like and exhausted CD8 TIL subsets through an unbiased analysis of single-cell heterogeneity. Moreover, we observed that among total CD8 TILs the most prominent subset corresponded to a transcriptomically distinct state that resembles that of effector memory T cells found in the context of acute infections.

### EM-like cells can be identified as Tcf1 *high* PD-1 *intermediate* CD8 TILs by flow cytometry

Our scRNA-seq data showed that the levels of *Tcf7* (encoding Tcf1) expression were high among naïve, memory-like and EM-like, and zero among exhausted cells (Figure 2 A). *Pdcd1* (encoding PD-1) expression was highest in exhausted and memory-like, intermediate in EM-like and absent in naïve CD8 TILs (Figure 2 A). To validate these observations at the protein level, we performed flow cytometry analysis of CD8 TILs infiltrating B16 tumors at day 12 post tumor engraftment. In agreement with previous studies (Thompson *et al.*, 2010) and consistently with our scRNA-seq data, we found both naïve (CD44^*low*^, 4 to 20%) and antigen-experienced (CD44^*high*^) cells among CD8 TILs. Next, CD44^*high*^ CD8 TILs were classified into four compartments according to Tcf1 and PD-1 expression: Tcf1^*low*^ PD-1^*high*^ (∼9% on average), Tcf1^*high*^ PD-1^*high*^ (∼31%) and Tcf1^*high*^ cells with low (∼21%) or intermediate (∼30%) PD-1 levels (Figure 2). According to the transcriptomic profiles and in line with previous evidence (Miller *et al.*, 2019; Siddiqui *et al.*, 2019), exhausted cells are mostly found in the Tcf1^*low*^ PD-1^*high*^ compartment and memory-like cells in the Tcf1^*high*^ PD-1^*high*^ gate. From our transcriptomics data EM-like cells were predicted to be in the Tcf1^*hig*^*h* PD-1^*int*^ compartment (Figure 2 A) and show over-expression of *Ly6c2, Cxcr3* and *Itgb7* (Figure 1 F). Indeed, the Tcf1^*high*^ PD-1^*int*^ compartment showed the highest levels of EM-like predicted markers Ly6C and CXCR3 and a higher proportion of ITGB7^+^ cells (Figure 2 B,C), confirming our predictions. Hence, EM-like cells can be identified by flow cytometry as Tcf1^*high*^ PD-1^*int*^ CD8 TILs.

**Figure 2.**
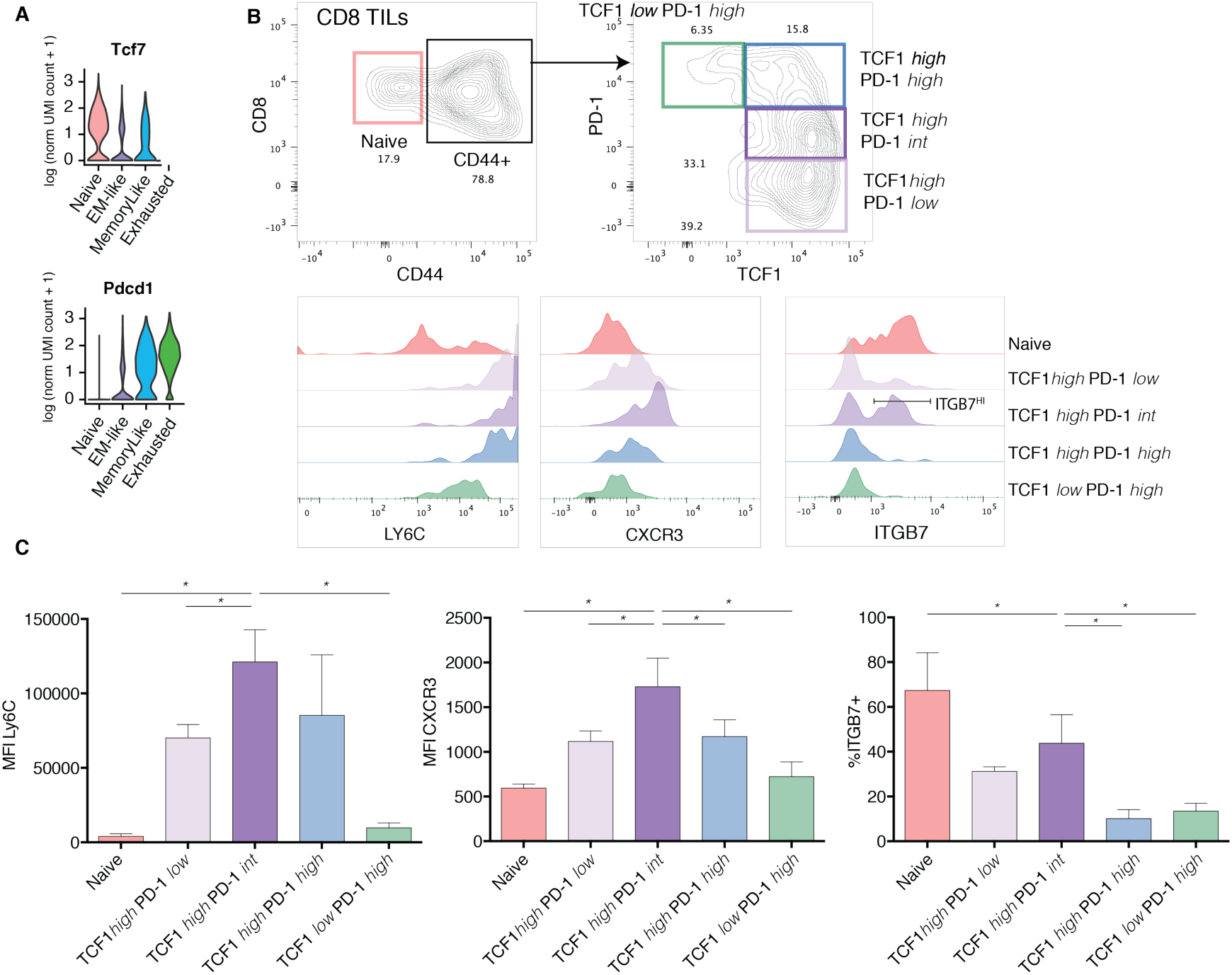
Flow cytometry validation of CD8 TIL populations. **A** Violin plots showing the decreasing and increasing expression levels (log-transformed normalized UMI counts +1) for *Tcf7* and *Pdcd1* for naïve, EM-like, memory-like and exhausted states. **B** Flow cytometry analysis of endogenous CD8 TILs from one representative tumor. Histograms show cell counts normalized by mode for naïve (CD44 *low*) T cells (red), Tcf1^*high*^ PD-1^*low*^ (light violet), Tcf1^*high*^ PD-1^*intermediate*^ (violet), Tcf1^*high*^ PD-1^*high*^ (blue) and Tcf1^*low*^ PD-1^*high*^ (green). **C** geometric Mean Fluorescence Index (MFI) for Ly6c and Cxcr3 and percentage of ITGB7^*high*^ cells for three tumors. Representative of two independent experiments. * denote statistically significant mean differences (Dunnett’s multiple comparisons test p-value < 0.05).

### Exhausted, Memory-like and EM-like CD8 TILs are clonally expanded and show partial clonal relatedness

In order to assess the clonal relatedness of CD8 TILs states, we analyzed TCR alpha and beta chain sequences in the >3500 single-cells shown in Figure 1 B (see Methods). We obtained full-length productive paired alpha and beta chains sequences (VJ or VDJ, respectively) in 81% of the CD8 TILs (Figure 3 A).

**Figure 3.**
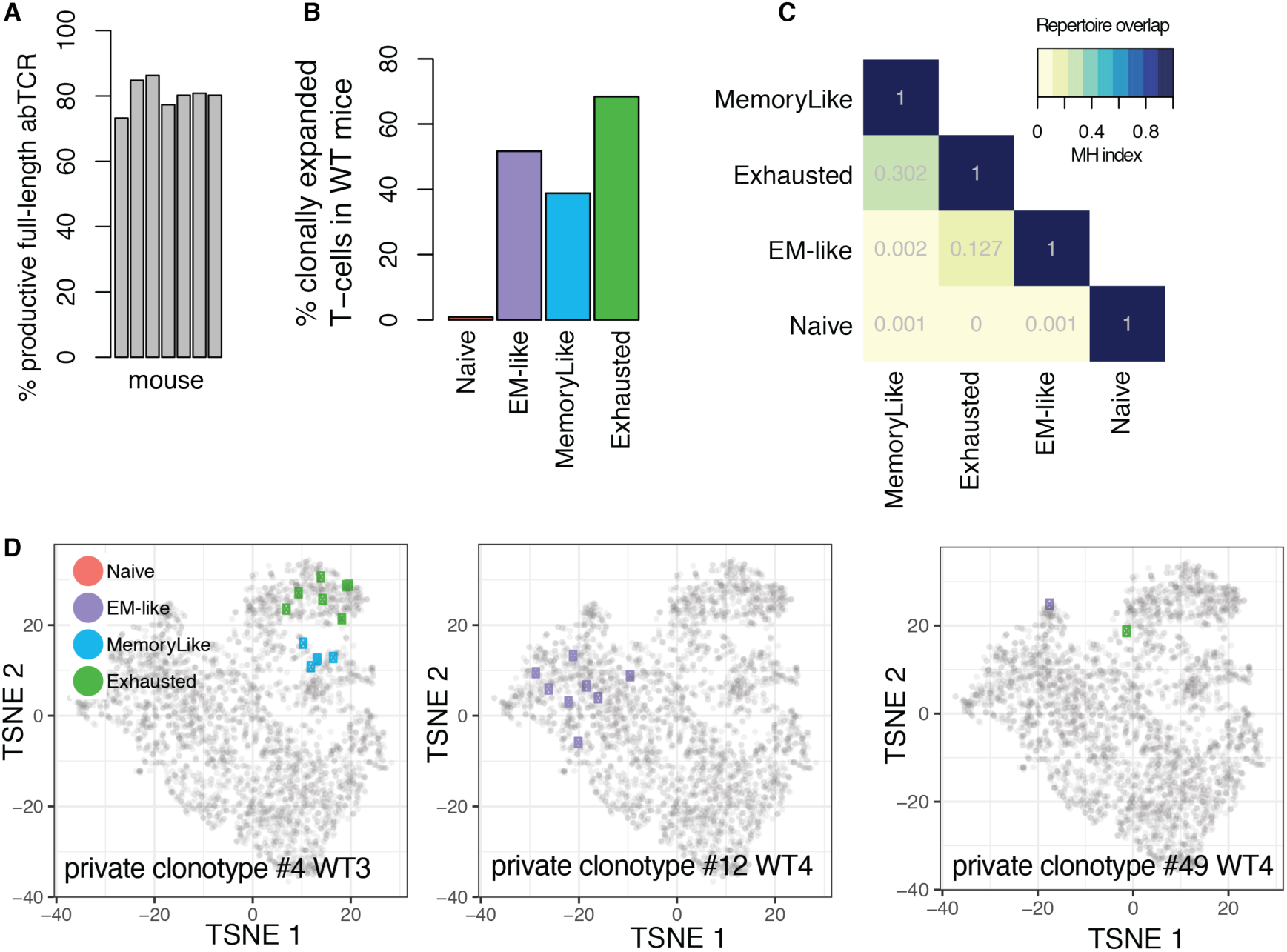
Clonal relatedness of CD8 TIL states. **A** Percentage of cells with productive TCR paired alpha/beta chains obtained in each of the 7 mice. **B** Percentage of T cells with non-unique (clonally expanded) TCR clonotype in each CD8 TIL cluster in endogenous responses in wild-type mice. **C** Repertoire overlap (Morisita-Horn index) between TIL states in wild-type mice. **D** Examples of clonally expanded T cells in wild-type mice. In each sub-panel, T cells with identical alpha and beta CDR3 sequences detected in individual mice are shown. Cell colors represent corresponding CD8 TIL states.

In each WT mouse, we identified T cells of the same clonotype, i.e. those expressing identical CDR3s for both alpha and beta T cell receptor chains. We found that between 10 and 39% of the TILs were expanded (i.e. their TCR were shared with at least another T cell in the same mouse). As expected, less than 1% of expanded T cells were found in the naïve state (Figure 3 B). In contrast, 68% of the T cells were expanded in the exhausted state, 52% in the EM-like state and 39% in the memory-like state. Expanded clonotypes did not match reported invariant chains and no known epitopes were found for these TCRs by literature and database searches. Furthermore, clonotypes in WT mice were largely private (mouse-specific) (see Methods and Supplemental Figure 5). As a control, a large clonal overlap was observed between different PMEL mice, due to the common transgenic PMEL TCR (Supplemental Figure 5).

Next, we evaluated TCR repertoire overlap between transcriptomic states. To this aim, we assessed TCR repertoire similarity using the Morisita-Horn (MH) similarity index that considers the relative frequencies of clonotypes between samples, where 0 indicates no overlap, and 1 is an exact match (Weinberger *et al.*, 2015). Interestingly, this analysis revealed that exhausted and memory-like states had a considerable repertoire overlap (MH index = 0.302, Figure 3 C), indicating the presence of shared T cell clones. A smaller overlap was observed between exhausted and EM-like states (MH = 0.127), indicating that these are also clonally related, yet to a lesser degree (examples of expanded clones are shown in Figure 3 D). Finally, very low overlaps were observed between EM-like and memory-like (MH = 0.002) or between naïve and any other state (MH = 0.001).

These results show that single clonotypes span both exhausted and memory-like states, in line with recent studies demonstrating that memory-like T cells can give rise to exhausted T cells in the tumor (Miller *et al.*, 2019; Siddiqui *et al.*, 2019). Furthermore, although EM-like and exhausted states presented largely distinct TCR repertoires, some clones were present in both, suggesting that occasionally EM-like cells (most likely tumor-specific) may yield exhausted T cells, yet to a lesser degree than the memory-like state.

### PD-1 checkpoint blockade expands EM-like cells

Multiple studies have established that PD-1 blockade expands intratumoral T cells and improves tumor control (Curran *et al.*, 2010; Wei *et al.*, 2017; Gubin *et al.*, 2018; Gide *et al.*, 2019). However, it is less clear how different CD8 T cell states are affected by immune-checkpoint blockade (ICB). Hence, here we aimed at evaluating the impact of ICB on the murine CD8 TIL transcriptomic landscape. To this end, we first extracted gene signatures of the exhausted, memory-like and EM-like states from our scRNA-seq dataset (Figure 4 A and Supplemental Table 1). Next, we performed gene-set enrichment analysis (GSEA) on published bulk RNA-seq data of CD8 TILs following anti-PD-1 therapy (raw data in murine sarcoma from (Gubin *et al.*, 2014). Our results indicated that ICB led to a selective enrichment of the EM-like signature (p=0.035, Figure 4 B). A similar effect was observed for non-small-cell lung cancer CD8 TILs upon ICB (Supplemental Figure 6 A, raw data from (Markowitz *et al.*, 2018)).

**Figure 4.**
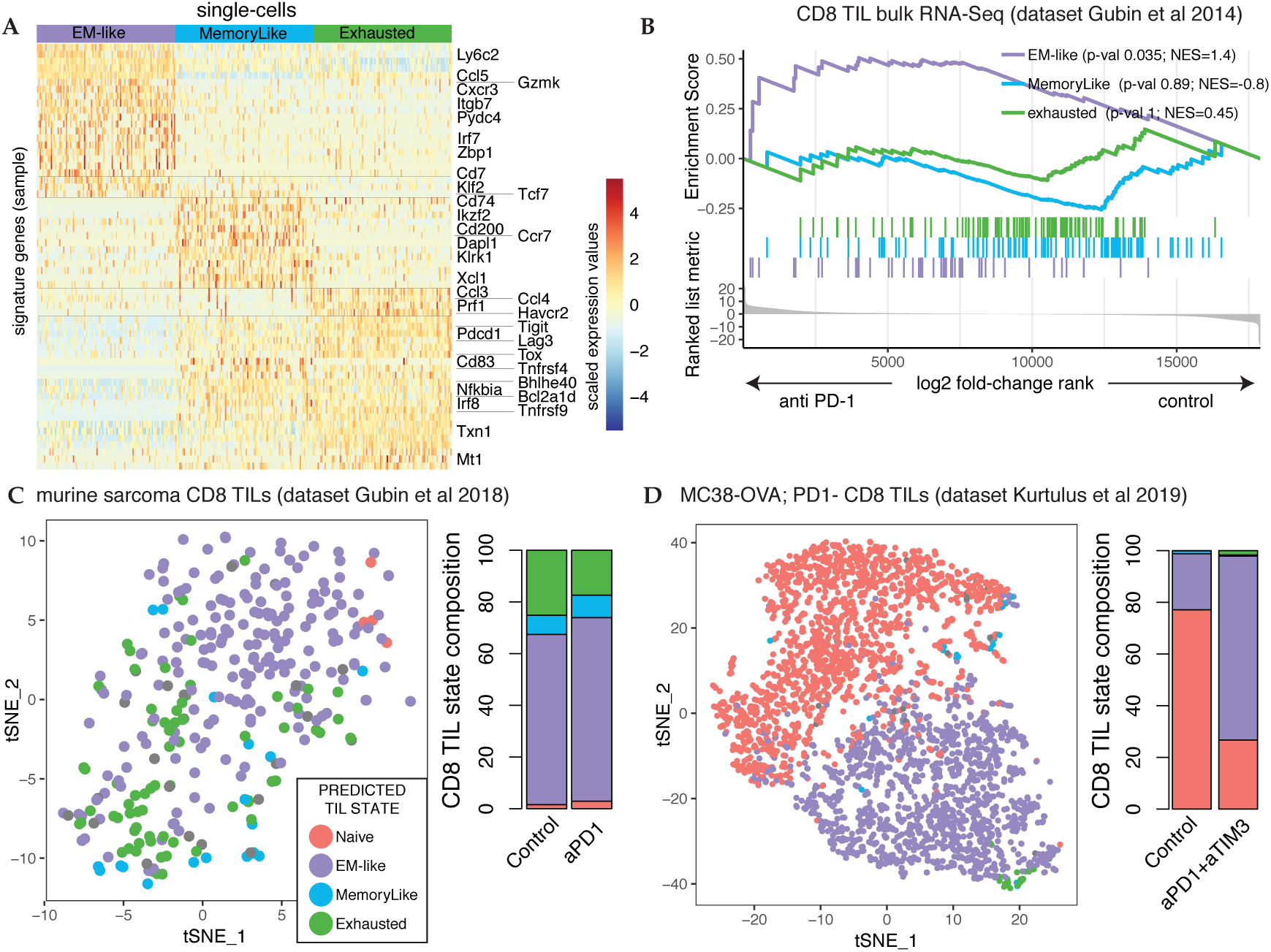
CD8 TIL state gene signature analysis of anti-PD-1 therapy. **A** Gene signatures of EM-like, memory-like and exhausted derived from our scRNA-seq analysis. For visualization, 100 random cells were sampled from each TIL state, and top differentially expressed genes are displayed. Horizonal lines separate groups of genes with similar expression patterns. **B** TIL state gene-signature enrichment analysis (GSEA) for the transcriptomic response of bulk endogenous CD8 TILs to PD-1 blockade (raw from (Gubin *et al.*, 2014)). NES=Normalized Enrichment Score. **C** tSNE plot of global transcriptomic similarity of CD8 TILs from murine sarcoma upon PD-1 blockade (dataset from (Gubin *et al.*, 2018)); **D** tSNE plot of global transcriptomic similarity of CD8 TILs from MC38 colon adenocarcinoma upon PD-1 and TIM3 blockade (dataset from (Kurtulus *et al.*, 2019), contains only PD-1 *low* TILs). Cell colors represent TIL state predictions by TILPRED.

In order to assess whether the bulk transcriptomic signature shift towards the EM-like state upon ICB is explained by differences in CD8 TIL states composition, we reanalyzed publicly available scRNA-seq data of CD8 TILs pre vs post ICB. To this aim, we developed a machine learning tool, named TILPRED, that predicts CD8 TIL states according to our definition (see Methods). TILPRED analysis of CD8 TIL scRNA-seq data from murine sarcoma ((Gubin *et al.*, 2018)) and MC38 colon adenocarcinoma (Kurtulus *et al.*, 2019) showed an increase in the proportion of EM-like TILs upon ICB at the single-cell level (Figure 4 C,D), consistent with the enrichment of the EM-like signature in the bulk transcriptomics data. Hence, our analyses indicate that EM-like T cells expand upon ICB.

Finally, we investigated whether ICB resulted in an enrichment of EM-like cells in cancer patients. Re-analysis of CD8 TIL scRNA-seq data from melanoma patients that responded to ICB (Sade-Feldman *et al.*, 2018) showed an increased proportion of predicted EM-like cells (p=0.032, see Methods and Supplemental Figure 7). These data suggest that intratumoral EM-like T cells in cancer patients might be modulated by ICB.

## DISCUSSION

In the past few years, single-cell transcriptomics and mass cytometry studies have revealed a large complexity of CD8 T cells in the tumor microenvironment, including multiple cellular states that have been referred to as exhausted, memory, naïve, effector, effector memory, etc., and this heterogeneity is likely to be a determining factor in therapy outcome (Tirosh *et al.*, 2016; Chevrier *et al.*, 2017; Kang *et al.*, 2017; Brummelman *et al.*, 2018; Guo *et al.*, 2018; Jerby-Arnon *et al.*, 2018; Sade-Feldman *et al.*, 2018; Zhang *et al.*, 2018; Li *et al.*, 2019; Yost *et al.*, 2019). However, as biological variability between patients and clinical samples is very large and tumor-specificity of T cells is rarely known, it is also crucial to define robust and unbiased tumor-infiltrating T cell transcriptomic states in murine models. Indeed, very recent studies in B16 murine melanoma model have shown that tumor-specific CD8 TILs can be found in the exhausted (PD-1^+^ TCF1^-^) and memory-like (“stem-like” PD-1^+^ TCF1^+^) states (Miller *et al.*, 2019; Siddiqui *et al.*, 2019).

Here we have defined a reference transcriptomic map of CD8 TILs in the common B16 melanoma model. Our single-cell RNA-seq analysis enabled us to robustly and unbiasedly define four distinct CD8 TIL transcriptomic states: exhausted, memory-like, naïve and effector memory-like (EM-like).

Consistently with previous observations, tumor-specific PMEL TILs were enriched in exhausted and memory-like states (Miller *et al.*, 2019; Siddiqui *et al.*, 2019). In contrast, total polyclonal CD8 TILs were enriched in the EM-like and naïve states. Therefore, an intriguing question is why the EM-like population is abundant in the endogenous polyclonal CD8 TIL compartment, but small among tumor-specific cells. A likely explanation is that the EM-like compartment is enriched in non tumor-specific cells, as multiple studies have shown the presence of large numbers of “bystander” T cells in human tumors (Simoni *et al.*, 2018; Scheper *et al.*, 2019). However, the presence of tumor-specific PMEL T cells in the EM-like state (Figure 1 G and Supplemental Figure 2) as well as the clonal expansion and partial TCR repertoire overlap between the EM-like and exhausted states (Figure 3 C,D) indicate that at least part of the EM-like population is indeed tumor-specific.

Compared to exhausted and memory-like, EM-like cells have lower expression of inhibitory receptors genes such as *Pdcd1, Tigit, Lag3* and of the transcription factor *Tox* (Supplemental Figure 6 D), which is essential to establish and maintain the epigenetic T cell exhaustion program enabling T cells to persist in the context of chronic antigenic stimulation (Alfei *et al.*, 2019; Khan *et al.*, 2019; Scott *et al.*, 2019; Yao *et al.*, 2019). Interestingly, reanalysis of published data indicated that tumor-specific CD8 TILs knock-out for *Tox* down-regulated the signature of the exhausted state and up-regulated the EM-like signature (and the memory-like signature to a lesser extent, Supplemental Figure 6 C). This suggests that *Tox*-KO tumor-specific CD8 TILs, which are unable to differentiate into exhausted cells, might remain in the pre-exhausted EM-like and memory-like states. Hence, our data suggest that the EM-like may be an early differentiation state of tumor-infiltrating CD8 TILs before receiving persistent antigenic stimulation. In this scenario, upon tumor migration, immunodominant clones in the EM-like state would rapidly activate the exhaustion program and differentiate in response to strong antigenic stimulation. In line with this hypothesis, our re-analysis of scRNA-seq data of tumor-specific CD8 TILs from Miller and colleagues (Miller *et al.*, 2019) indicated that while 12% of EM-like cells were detected at day 10 (Supplemental Figure 4), only 1% were detected at day 20 post-tumor engraftment, suggesting a temporal progression towards more differentiated states. Cognate antigen dose and T cell avidity might be relevant factors controlling the dynamics of such intratumoral differentiation. For example, it has been shown that increased antigen dose and T cell avidity promote CD8 T cell differentiation into the exhausted phenotype in cancer and chronic infection (D. T. Utzschneider *et al.*, 2016; Martínez-Usatorre *et al.*, 2018). It is possible that subdominant clones, less subject to persistent stimulation, undergo a retarded differentiation and accumulate in the EM-like state. Hence, our results have potential implications on the understanding of intratumoral differentiation of CD8 T cells, an open central question in immunology (Blank *et al.*, 2019).

Our flow cytometry analysis revealed that the EM-like population is enriched within the Tcf1^*high*^ PD-1^*intermediate*^ CD8 TIL compartment and expresses high levels of the surface markers Ly6C, CXCR3 and ITGB7, as predicted from the transcriptomic analysis. These are novel EM-like CD8 TILs surface markers that can be useful in functional studies of this population. Ly6C is an adhesion molecule expressed by neutrophils, monocytes, dendritic cells and also in subsets of CD4 and CD8 T cells, including memory CD8 T cells (Walunas *et al.*, 1995; Pihlgren, 1996; Cerwenka *et al.*, 1998; Lee *et al.*, 2013). Interestingly, Ly6C^+^ CD8 T cells with effector memory phenotype isolated from spleens of tumor-primed mice have shown anti-tumor activity *in vitro* (Piranlioglu *et al.*, 2019), supporting the hypothesis that tumor-specific EM-like cells are related to pre-exhaustion states. The chemokine receptor CXCR3 is important for the recruitment of T cells into inflamed peripheral tissue (Groom and Luster, 2011) and is also required by CD8 TILs for effective response to anti-PD-1 therapy (Chow *et al.*, 2019). As the EM-like state expressed the highest levels of CXCR3, this chemokine system might contribute to the enrichment of EM-like cells upon anti-PD-1 therapy observed in our study. The EM-like state -together with naïve cells-also showed differential expression of the integrin subunit β7 (*Itgb7*). At the protein level however, ITGB7 was expressed only by a subset of EM-like cells (Figure 2 C). Of note, *Itgb7* was co-expressed in EM-like cells with the integrin subunit α4 (*Itga4*) but not with *Itgae* (CD103), with which ITGB7 dimerizes in tissue-resident T cells (Corgnac *et al.*, 2018). Instead, the α4β7 integrin has been previously shown to mediate lymphocyte migration to gut-associated lymphoid tissue and might have a different function in the context of tumors (Denucci, Mitchell and Shimizu, 2009).

Multiple studies have established that PD-1 blockade increases T cell infiltrations in tumors leading to improved anti-tumor control (Curran *et al.*, 2010; Wei *et al.*, 2017; Gubin *et al.*, 2018; Gide *et al.*, 2019). Moreover, recent studies have shown that ICB promotes expansion of tumor-specific memory-like cells and their differentiation into (terminally) exhausted cells (Miller *et al.*, 2019; Siddiqui *et al.*, 2019). However, it is less clear how the CD8 TIL landscape is impacted by ICB.

Our meta-analysis of bulk and single-cell transcriptomic data showed a selective enrichment of the EM-like signature (e.g. *Cxcr3, Ly6c2, Ccl5, Gzmk, Itgb7*) following ICB. As the EM-like is a relatively undifferentiated state, we reasoned that the EM-like signature would not be enriched upon ICB among T cells subsets that were already differentiated before treatment. Indeed, we observed that *in vitro* activated tumor-specific cells did not up-regulate the EM-like signature following adoptive transfer and ICB but instead up-regulated the exhaustion signature (e.g. *Mt1, Havcr2, Prf1, Gzmb*, etc., Supplemental Figure 6 B, sequencing data from (Mognol *et al.*, 2017)). This suggests that the enrichment in EM-like cells upon ICB depends on the expansion of relatively undifferentiated clones whereas tumor-specific T cells that have already activated the exhaustion program will only progress further towards exhaustion in response to ICB. In line with our observations, other studies have shown up-regulation of *Cxcr3, Ccl5* and *Ifit3* (EM-like state genes) or expansion of PD-1^*low*^ CD8 T cells (that partially contain EM-like cells, Figure 4 D) among CD8 TILs following ICB in murine models (Gubin *et al.*, 2018; Kurtulus *et al.*, 2019). In humans, a CD8 TIL subset differentially expressing GZMK and CXCR3 (EM-like state markers) has been previously observed in scRNA-seq analysis of basal and squamous cell carcinoma (Yost *et al.*, 2019), hepatocellular carcinoma (Kang *et al.*, 2017) and small-cell lung cancer (Guo *et al.*, 2018), suggesting that a similar population might be conserved between mice and human. However, in these studies tumor specificity was unknown, limiting our understanding of EM-like TILs in cancer patients. Our results will facilitate future investigations aimed at defining the function of intratumoral EM-like cells, which may provide novel markers of T cell response to immune checkpoint blockade.

In conclusion, our scRNA-seq study defined a reference map of the transcriptomic landscape of CD8 TILs in B16 melanoma. As such, this resource provides a base upon which to interpret more complex CD8 TIL transcriptomic landscapes such as those derived from clinical samples, where T cell specificity and clonality are usually unknown. Our CD8 TIL state predictor is available as an R package at https://github.com/carmonalab/TILPRED and a streamlined web interface for exploring our scRNA-seq data is available at https://tilatlas.shinyapps.io/B16_CD8TIL_10X/.

## MATERIALS AND METHODS

### Mice

For scRNA-seq 6-12 week-old male C57BL/6 mice (CD45.1^+^) and PMEL (Jackson Laboratory, Cat#005023) were bred and housed under SPF conditions in the conventional animal facility of the University of Lausanne. Experiments were performed in compliance with the University of Lausanne Institutional regulations and were approved by the veterinarian authorities of the Canton de Vaud.

### Tumor experiments and isolation of TILs

Right flanks of mice (C57BL/6 or PMEL) were shaved and B16F10 cells (2.5×10^5^) were injected subcutaneously. Tumor volumes were estimated by measuring the tumor size in two dimensions using a caliper. The tumor volume was calculated according to the formula (length x× width^2^)/2. Mice were sacrificed at the indicated time point and the weight of the excised tumor mass was determined.

Tumors were excised post 15 days of tumor engraftment, manually dissociated and digested enzymatically with Tumor Dissociation Kit (Miltenyi Biotec). Digested tumors were mashed through 70 µm filters. Hematopoietic cells were further purified using a discontinuous Percoll gradient (GE Healthcare). Cells at the interface were harvested and washed twice before further use. For cell sorting, tumor infiltrating T cells were further purified using mouse CD8 T cell enrichment kit (StemCell Technologies) and sorted by flow cytometry using following antibodies - CD8a (Clone 53-6.7; eBiosciences), CD45 (Clone MI/89; eBiosciences) and Zombie Aqua (423102; Biolegend). The purity of sorted cells was greater than 99%. Flow sorted tumor infiltrating CD8 T cells were thus used for single cell RNA sequence purpose.

### 10x Genomics single-cell gene expression sample processing and sequencing

After sorting, intratumoral CD8 T cells were loaded into a Chromium Single Cell Instrument (10x Genomics, Pleasanton, CA) together with beads, master mix reagents (containing RT enzyme and poly-dt RT primers) and oil to generate single-cell-containing Gel Beads-in-emulsion (GEMs). Single-cell Gene Expression libraries were then prepared using Chromium Single Cell 5’ Library & Gel Bead Kit (PN-1000006) following the manufacturer’s instruction (protocol CG000086 Rev E). With this procedure, the cDNAs from distinct droplets harbor a distinct and unique 10x “cell barcode”. These sequencing libraries were loaded onto Illumina NextSeq High Output Flow Cells and sequenced using read lengths of 26 nt for read1 and 132 nt for read2. The Cell Ranger Single Cell Software Suite 2.1.1 (https://support.10xgenomics.com/single-cell-gene-expression/software/pipelines/latest/using/mkfastq) was used to perform sample demultiplexing, barcode processing, reads downsampling per sample (down to 118,806 mean reads per cell) and single cell 5’ gene counting using default parameters and mouse genome assembly mm10.

### Single-cell TCR sequencing and analysis

Intratumoral CD8 T cells cDNAs that were generated as an intermediate step during the aforementioned single-cell gene expression libraries preparation were subsequently used in a distinct workflow. Briefly, the 10x mouse V(D)J Enrichment Kit (PN-1000071) was used to enrich for TCR sequences, after which V(D)J-enriched libraries were constructed with the Chromium Single Cell 5’ Library Construction Kit (PN-1000020). The Cell Ranger Single Cell Software Suite 2.2.0 (https://support.10xgenomics.com/single-cell-gene-expression/software/pipelines/latest/using/mkfastq) was used to perform sample demultiplexing, barcode processing and clonotype identification, using default parameters.

For each mouse separately, T cells sharing identical alpha and beta TCR chains were assigned to a unique clonotypes ID. PMEL-specific cells were identified based on expression of the transgenic PMEL-specific TCR (CASSFHRDYNSPLYF and CAVNTGNYKYVF for beta and alpha CDR3s, respectively, (Overwijk *et al.*, 2003), GenBank entries EF154513.1 and EF154514.1, Supplemental Figure 6 B). Single-cell TCR sequences are available as Supplementary Files at GEO under accession GSE116390).

TCR repertoire similarity between mice (Supplemental Figure 5 A) and TIL states (Figure 3 B) was calculated using the Morisita-Horn (MH) index as implemented in the R package ‘divo’ (Rempala and Seweryn, 2013).

### Processing, dimensionality reduction and clustering of newly generated scRNA-seq data

A total of 7174 single-cell transcriptomes were obtained from 7 samples (4 WT + 3 PMEL mice) after Cell Ranger pre-processing and UMI quantification. Next, UMI counts were normalized by dividing them by the total UMI counts in each cell and multiplying by a factor of 10,000. Then, we took the log of the normalized UMI counts prior sum of 1 (log normalized UMI counts+1). Then, cells were filtered based on quality control and expression of CD8 T cell markers to remove low-quality cells, doublets and contaminating non CD8 T cells, as follows. Upon examination of parameters distributions, we filtered out cells having less than 1,500 or more than 30,000 UMIs; cells having less than 1,500 or more than 5,000 detected genes, cells in which ribosomal protein coding genes represented more than 50% of UMI content and cells in which mitochondrial protein coding genes represented more than 5% of UMI content. The 5542 quality-passed cells were further filtered based on expression of CD8 T cells markers: 3574 cells expressing *Cd8a, Cd8b1* and *Cd2* but not *Cd4* were kept for further analysis (processed data available as supplementary file in GEO entry).

For dimensionality reduction, we first identified the set of most variable genes using Seurat 2.3.4 method ‘mean.var.plot’ (using 20 bins, minimum mean expression = 0.25 and z-score threshold for dispersion = 0), which identified 1107 highly variable genes while controlling for the relationship between variability and average expression. Briefly, this method divides genes into 20 bins based on average expression, and then calculates z-scores for dispersion (calculated as log(variance/mean)) within each bin. From this initial set of highly variable genes, we removed 204 genes involved in cell cycle (as annotated by Gene Ontology under code GO:0007049 or highly correlated with them, i.e. with Pearson’s correlation coefficient > 0.5) or coding for ribosomal or mitochondrial proteins. The remaining 903 highly variable genes were used for dimensionality reduction using Principal Components Analysis (PCA). PCA was performed on standardized gene expression values by subtracting from normalized UMI counts, their mean and dividing by the standard deviation. Upon scree plot inspection of PCA eigenvalues contributions, we selected the first 10 Principal Components for clustering and tSNE visualization. For visualization, we used tSNE with default parameters (perplexity = 30 and seed set to 12345). For clustering, we performed hierarchical clustering on the top 10 PCs using Euclidean distance and Ward’s criteria. Silhouette coefficient analysis over different number K of clusters indicated a big drop of cluster silhouette after K=4, and this was selected as the optimal number of clusters. To evaluate clustering robustness, we additionally run K-means (with K=4) and the shared nearest neighbor (SNN) modularity optimization clustering algorithm implemented in Seurat 2.3.4 with resolution parameter = 0.3 (which produced 4 clusters) and other parameters by default. Clustering agreement analysis using adjusted Rand Index (as implemented in mclust R package (Scrucca *et al.*, 2016)), indicated high agreement between the three clustering results (Rand Index 0.70-0.81). Moreover, this analysis indicated that the SNN clustering produced the most consistent result with the other two (with Rand Index of 0.81 against hierarchical and 0.76 against K-means, while K-means vs hierarchical had 0.7), and therefore was kept as the final clustering solution.

### Gene Signature analysis

To obtain cluster-specific gene signatures, we performed differential expression analysis of each cluster against the others using MAST (Finak *et al.*, 2015) with default parameters, and further required that for each cluster, differentially expressed genes had a log fold-change higher than 0.25, were expressed at least in 10% of its cells, and that this fraction is at least 10% higher than in the other clusters. Lists of differentially expressed genes in each cluster can be found in Supplemental Table 1.

To identify cycling cells we evaluated enrichment of the cell cycle signature (Supplemental Table 2) in each cell, using the Area Under the Curve (AUC) method implemented in AUCell Bioconductor’s package (Aibar *et al.*, 2017). Briefly, the AUC value represents the fraction of genes, within the top 1500 genes in the ranking (ordered by decreasing expression) that are included in the signature. Upon examination of AUCs distribution, we set an AUC cut-off of 0.2 for cell cycling classification (which corresponded to a z-score of ∼2.5).

For CD8 T cell subtypes reference signature enrichment analysis, we first collected different T cell signatures from literature and generated other signatures by performing differential expression analysis (using Geo2R NCBI Gene Expression Omnibus tool, with default parameters) from public transcriptomic datasets. Cut-off values for log fold-change (between 1 and 3) and adjusted p-value (either 0.05 or 0.01) were set in order to obtain gene sets of similar size (in the order of a hundred genes), as described below. The list of genes in each signature can be found in Supplemental Table 2.

For signatures of naïve (CD44-), effector (KLRG1^+^ at day 4.5 p.i.) and memory (day 60 p.i.) virus-specific CD8 T cell isolated in the context of acute infection (“Memory vs effector (acute inf.)”), we used the LCMV Armstrong virus infection data from (Sarkar *et al.*, 2008), GEO accession GSE10239, considering differentially expressed genes of effector vs naive, memory vs naïve and effector vs memory. For every gene signature, up to 200 top genes were kept sorted by fold-change from the differentially expressed genes (defined as FDR adjusted p-value < 0.05 and |log2 fold-change| > 1). The signatures of (PD-1^+^ Tcf1^+^) memory-like vs (PD-1^+^ Tcf1^-^) exhausted virus-specific (P14) CD8 T cells in LCMV chronic infection clone 13 (“memory-like vs exhausted (chronic inf.)”) were obtained from ((Utzschneider *et al.*, 2016), GSE83978) by taking differentially expressed genes between PD-1^+^ *Tcf7*:GFP^+^ vs PD-1^+^ *Tcf7*:GFP^-^. The signatures of (PD-1^+^ Tcf1^+^) memory-like vs (PD-1^+^ Tcf1^-^) exhausted tumor-specific(P14) CD8 T cells in B16-gp33 melanoma tumors (“memory-like vs exhausted (tumor)”) were obtained from GEO GSE114631 (Siddiqui *et al.*, 2019) by taking differentially expressed genes between *Tcf7*:GFP^+^ vs *Tcf7*:GFP^-^. For the signatures of tumor-infiltrating vs spleen-derived in acute infection at day 8 post listeria infection (“Tumor vs acute infection”), we used the data of (Schietinger *et al.*, 2016) (GEO GSE60501).

For cell cycle signature we used the gene set (cellCycle_union, Supplemental Table 2) obtained by combining the G1/S and G2/M signature genes from (Tirosh *et al.*, 2016) (cellCycle, Supplemental Table 2) with the set of genes whose expression in our dataset were highly correlated with cell-cycle related genes (Pearson’s correlation > 0.5, cellCycleCorrelated, Supplemental Table 2).

To evaluate associations (overlap) between cluster signatures and reference signatures (Figure 1 E and Supplemental Figure 1), for each pair of cell cluster signature genes and reference signatures, contingency tables were calculated by counting how many genes among all expressed genes (15337) are present or absent in the cluster or the reference signature, and then one-sided Fisher’s exact test with FDR adjustment for multiple testing were used to calculate statistical significance of associations.

For the quantification of stemness (*Tcf7, Lef1, Sell, Il7r*), cytotoxicity (*Prf1, Fasl, Gzmb*) and inhibition/exhaustion (*Pdcd1, Havcr2, Tigit, Lag3, Ctla4*) scores, we calculated single-cell AUC enrichment scores (using AUCell package, as described for cell cycling). For Figure 1 D, average enrichment scores for each cluster (after removing cycling cells from all clusters) were scaled to range between 0 and 1 and plotted using ggradar R’s package (https://github.com/ricardo-bion/ggradar).

To evaluate enrichment of EM-like, memory-like and exhausted signatures in public bulk RNA-seq data of CD8 T cells upon PD-1 blockade or TOX KO, we performed signature enrichment analysis (GSEA) using Bioconductor package clusterProfiler with default parameters (version 3.10.1, (Yu *et al.*, 2012)). Reads were pre-filtered using Trimmomatic v0.39 (Bolger, Lohse and Usadel, 2014); transcript abundances were estimated using Salmon v0.14.0 (Patro *et al.*, 2017) with default parameters and mouse reference transcriptome version GRCm38, summarized at gene-level using tximport v1.10.1 (Soneson, Love and Robinson, 2015) and differential gene expression analysis was conducted using DESeq2 v1.22.2 with default parameters (Love, Huber and Anders, 2014). RNA-seq data of CD8 TILs in murine sarcoma (tumors were harvested at day 12 post-transplant, 3 days after treatment) was obtained from GEO accession GSE62771 (anti-PD-1 vs control) (Gubin *et al.*, 2014). RNA-seq data of CD8 TILs in non-small cell lung cancer (anti-PD1 vs control) were obtained from GEO accession GSE114300 (Markowitz *et al.*, 2018). RNA-seq data from adoptively transferred (*in vitro* activated) OT-1 cells infiltrating B16-OVA tumors (anti-PD1 vs control) were obtained from GEO GSE93014 (Mognol *et al.*, 2017). Differential expression data from tumor-specific CD8 TILs (liver cancer) knockout for TOX (TOX-KO) vs wildtype were obtained from Supplementary Table 1 of (Scott *et al.*, 2019).

### Cluster validation using independent dataset

To confirm the robustness of the transcriptomic states identified in our data, we reanalyzed an independent publicly available single-cell RNA-seq dataset of ∼400 CD8 TILs from B16 melanoma tumors (Singer et al 2016). Gene expression data processing and clustering was conducted in the same way as described for our dataset. Consistently, four distinct transcriptomic states were identified corresponding to naïve, terminal exhausted, memory-like and EM-like populations (Supplemental Figure 3 A, B), plus a cluster of cycling cells in proximity to exhausted and memory-like cells. Despite the different technologies used (smart-seq2 vs 10X 5’ counting) we found a remarkable correspondence between the CD8 TIL states identified in the two datasets, both in terms of gene markers (Supplemental Figure 3 B) and systematic comparison of gene signatures (Supplemental Figure 3 G). To evaluate associations (overlap) between cluster signatures from the two datasets, one-sided Fisher’s exacts test with FDR adjustment for multiple testing were used (Supplemental Figure 3 G).

### Flow Cytometry Analysis of CD8 TILs

For the flow cytometry analysis purified CD8 cells specific for the surrogate tumor antigen LCMV gp33 (P14) were adoptively transferred (10^6^ cells) into naive C57BL/6 (B6) mice (CD45.1). One day later, mice were implanted subcutaneously with B16 melanoma cells expressing LCMV gp33 (B16-gp33). Tumors were excised post 12 days of tumor engraftment and tumor infiltrating T cells were isolated as described above. Surface staining was performed with mAbs for 30 min at 4°C in PBS supplemented with 2% FCS and 0.01% azide (FACS buffer) using antibodies listed in table below.

For intranuclear staining cells were first stained at the surface before fixation and permeabilization using the transcription factor staining kit (eBioscience) followed by intranuclear staining for transcription factor.

Flow cytometry measurements of cells were performed on an LSR-II or Fortessa flow cytometer (BD). Data were analyzed using FlowJo (TreeStar).

| ANTIBODIES | SOURCE | IDENTIFIER |
| --- | --- | --- |
| Anti-Mouse CD8a – APC-Cy7 | eBioscience | Clone 53-6.7 |
| Anti-Mouse CD45.1 – AF680 | eBioscience | Clone A20 |
| Anti-Mouse CD45.2 – PerCP Cy5.5 | eBioscience | Clone 104.2 |
| Anti-Mouse PD1 – PE Cy7 | Biolegend | Clone RMP1-30 |
| Anti-Mouse CD45R/B220 – BV 711 | Biolegend | Clone RA3-6B1 |
| Anti-Mouse Ly-6C – BV 785 | Biolegend | Clone HK1.4 |
| Anti-Mouse CD183 (CXCR3) – BV 650 | Biolegend | Clone CXCR3-173 |
| Anti-Mouse CD44 – BV 605 | Biolegend | Clone IM7 |
| Anti-Human/Mouse integrin b7 (ITGB7) - FITC | Biolegend | Clone FIB504 |
| Mouse anti-rabbit IgG - PE | eBioscience | polyclonal |
| Rabbit anti-mouse Tcf1 | Cell signaling | C63D9 |

### Construction and validation of the Tumor-infiltrating CD8 T cell transcriptomic state predictor (TILPRED)

Integrated knowledge on CD8 tumor-infiltrating T cells states was used to develop a novel transcriptomic classifier of CD8 T cell states for mice and human (TILPRED). A major challenge for the classification of single-cell transcriptomes across experiments is that gene expression measurements are affected by the large variability in single cell library sizes, and experimental and data processing protocols used. To overcome some of these issues, our classifier uses only gene expression rankings to quantify TIL state signature enrichment. Moreover, since cell cycle has a significant transcriptomic impact (McDavid *et al.*, 2014; Scialdone *et al.*, 2015; Barron and Li, 2016), we also included cell cycle signature detection in TILPRED, which allows for quantification of cycling T cells in the different states.

To classify tumor-infiltrating CD8 T cells from single-cell RNA-seq data we first generated 12 sets of differently expressed genes (DEG) corresponding to all possible pairs between the four transcriptomic states. Differentially expression between pairs of clusters was assessed with MAST 1.8.2 using default parameters (after excluding cycling cells in all cluster, in order to avoid this confounding factor), and further filtering genes with a log fold-change higher than 0.5, expressed at least in 10% of one of the clusters, and with a cellular detection rate difference of at least 10% between clusters. Next, pairwise DEG sets were further filtered to keep those having orthologs in humans (as annotated in Ensembl 95 (Zerbino *et al.*, 2018), or NCBI HomoloGene 68 if orthologs missing in Ensembl).

The 12 pairwise DEG sets were then used to score all single-cells in our dataset, calculating gene set enrichment AUC (area under the ‘enrichment’ curve) within the top 1500 genes in the ranking, using AUCell (Aibar *et al.*, 2017). Next, the AUC scores for the 12 pairwise DEG (a training set of 70% of the cells) were used to train a multinomial logistic regression model with lasso penalty using R package *glmnet* (Friedman, Hastie and Tibshirani, 2010) with parameter alpha = 1 (lasso penalty) and intercept set to 0. The regularization parameter lambda for the lasso was evaluated by 10-fold cross-validation using cv.glmnet function, and was set to the minimum value (lambda = e^-^6) for which the mean cross-validated error (cve) was at most 50% higher (i.e. cve = 0.539) than the minimum mean cross validation error (cve = 0.374, for lamda = 4.2 e^-^5).

Performance of the model was assessed in the test set (remaining 30% of the cells) assigning T cell state predictions to all cells where the model output (logistic response variable) was higher than 0.5 in any state (∼5% of the cells did not pass the threshold and were annotated as ‘unknown’); its accuracy was 0.94 (95% CI: 0.92, 0.95) with a mean specificity of 0.98 and a mean sensitivity of 0.90. Finally, the model was trained on the full dataset and implemented as an R package, publicly available at http://github.com/carmonalab/TILPRED.

To evaluate predictive performance in an independent dataset, we compared TILPRED predictions against the result of unsupervised clustering of the CD8 TIL dataset of (Singer *et al.*, 2016) (GEO GSE86042). The clustering analysis was performed in the same way as for our dataset (on the first 10 principal components of the most highly variable genes), except that instead of using a unique clustering solution (such as SNN clustering used to defined clusters in our dataset), we used as T cell states ‘ground truth’ the consensus between SNN, K-means and hierarchical clustering (using Ward’s criteria) solutions. Classification accuracy using this dataset was 91%.

### TILPRED analysis of public datasets

For prediction of CD8 TIL states in the MC38 dataset (NCBI GEO GSE122969 (Kurtulus *et al.*, 2019)), high-quality cells were filtered based on number of detected genes (between 500 and 5000), number of UMIs (1K to 50K) and percentage of UMIs mapping to mitochondrial (<10%) or ribosomal genes (<60%). CD8 T cells were further filtered by based on co-expression of *Cd2, Cd8a, Cd8b1* and *Cd3g* (>= 1 UMI each) and lack of *Cd4* expression. For the sarcoma dataset (NCBI GEO GSE122969 (Gubin *et al.*, 2018)), after examining distributions, high-quality cells were filtered based on number of detected genes (between 500 and 5000), number of UMIs (1.5K to 20K) and percentage of UMIs mapping to mitochondrial (<10%) or ribosomal genes (<50%). CD8 T cells were further filtered from other immune cell infiltrates based on co-expression of *Cd2, Cd8a* and *Cd8b1* (>= 1 UMIs each), lack of *Cd4* expression (0 UMI) and lack of -or low expression- of *Fcer1g* and *Tyrobp* (<2 UMIs). For both filtered datasts, TILPRED was run with default parameters.

For the prediciton of CD8 TIL states in the melanoma patients dataset (GEO GSE120575 (Sade-Feldman *et al.*, 2018)), high-quality cells were filtered based on number of detected genes (between 1000 and 6000) and percentage of UMIs mapping to ribosomal genes (<10%). CD8 T cells were further filtered from other immune cell infiltrates based on co-expression of CD2, CD8A, CD8B (>= 1 UMIs each), lack of CD4 expression (0 UMI) and lack of -or low expression-of FCER1G, TYROBP, SPI1, IGKC, IGJ, IGHG3 (<7 UMIs). Next, TILPRED was run using parameters set for human genes (human=TRUE) and using with lower score threshold for prediction (scoreThreshold=0.3, instead of the default 0.5), in order to increase sensitivity to weaker signals (as TILPRED was trained for mouse data). For the visualization of these data in Figure 3, we used Seurat 2.3.4 to identifiy highly variable genes and perform dimensionality reduction using tSNE on the first 10 Principal Components computed on centered and scaled normalized expression values (log (UMI counts / sum UMI count in cell * 10,000 + 1)). The four CD8 TIL states (naïve, EM-like, exhausted and memory-like) as well as cycling TILs were predicted in all patients (Supplemental Figure 7A). To evaluate compositional shifts upon therapy, we selected patients having samples before and after ICB and at least 30 cells in each sample (Supplemental Figure 7B). Seven patients matched these criteria, 3 of which responded (P1, P7, P28) and 4 that failed to respond to ICB (P2, P3, P12, P20). Interestingly, compared to nonresponders, responders had an increased proportion of EM-like cells upon therapy (Supplemental Figure 7 C, p=0.0317 one-sided Wilcoxon test).

For the prediction of TIL states of the Miller dataset of (B16-OVA) tumor-specific tumor-infiltrating CD8^+^ T cells, UMI counts matrix was obtained from GEO (accession GSE122675). Upon examination, high-quality CD8 T cells were filtered as those having between 1000 and 5000 expressed genes, 1000 and 30,000 UMI counts, less than 6% of UMI counts mapping to mitochondrial genes, less than 50% of genes mapping to ribosomal proteins coding genes, and that expressed *Cd2, Cd3g, Cd8a, Cd8b1* and did not express *Cd4* or high levels of *Tyrobp* or *Spi1* (that are associated to CD4 T cells or myeloid cells, respectively). Next, cycling cells were further filtered (those with an AUC score higher than 0.1 for the ‘cell cycle’ signature) and cell states were predicted using TILPRED with default parameters.

### WebApp deployment for scRNA-seq data exploration

Our web application uses iSEE (interactive SummarizedExperiment Explorer) (Rue-Albrecht *et al.*, 2018) and R Shiny (https://shiny.rstudio.com), and is available at https://tilatlas.shinyapps.io/B16_CD8TIL_10X/.

## SUPPLEMENTAL MATERIAL

Supplemental Table 1 (SupplementalTable1.csv): Exhausted, Memory-like and EM-like gene signatures

Supplemental Table 2 (SupplementalTable2.csv): Reference gene signatures used for signature enrichment analyses

Supplemental Material. Contains Supplemental Figures.

## DECLARATIONS

### ETHICS APPROVAL

Animal experiments were performed in compliance with the Helsinki Declaration and the University of Lausanne Institutional regulations, and were approved by the veterinarian authorities of the Canton de Vaud.

### CONSENT FOR PUBLICATION

All authors have approved the manuscript.

### DATA AVAILABILITY

The single-cell sequencing data generated in this study are available in NCBI Gene Expression Omnibus (GEO) under accession number GSE116390. Single-cell datasets re-analysed in this study are available in NCBI GEO under accession numbers GSE86042 (CD8 TILs from B16 melanoma, (Singer *et al.*, 2016)), GSE122969 (PD-1-CD8 TILs from MC38, (Kurtulus *et al.*, 2019)), GSE119352 (whole-tumor sarcoma, (Gubin *et al.*, 2018)), GSE120575 (CD8 TILs of melanoma patients, (Sade-Feldman *et al.*, 2018)).

### AUTHORS’ CONTRIBUTIONS

SJC, WH and DG conceived and designed the project, SJC analyzed the data and developed the bioinformatics tools; IS performed wet experiments and analyzed data; MB contributed to data analysis; SJC, WH and DG wrote the manuscript. All authors read and approved the final manuscript.

## ACKNOWLEDGMENTS

We are grateful to Amaia Martinez Usatorre, Jesus Corria Osorio and Julien Racle for fruitful discussions and critical reading of the manuscript; to Prof. Pedro Romero for kindly providing the PMEL mice; to the Gene Expression Facility of EPFL (École Polytechnique Fédérale de Lausanne) that performed the 10x Genomics single-cell assays and the UNIL Flow Cytometry Facility for their assistance. Computations were performed at the Vital-IT Center for high-performance computing of the Swiss Institute of Bioinformatics (http://www.vital-it.ch).

