## Supplemental Material for "Deciphering the transcriptomic landscape of tumor-infiltrating CD8 lymphocytes in B16 melanoma tumors with single-cell RNA-Seq"

Legend Supplemental Table 1 – Reference Gene signatures

Legend Supplemental Table 2 – Differentially expressed genes between TIL clusters

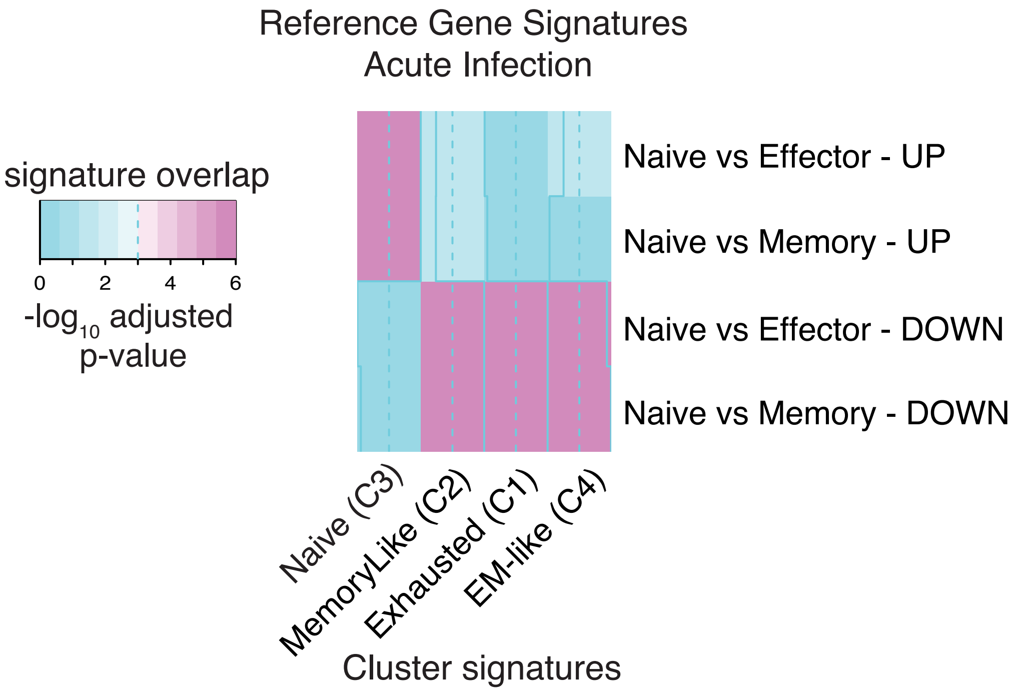


**Supplemental Figure 1** Gene signature enrichment analysis against reference CD8 T-cell subtypes signatures observed in acute infection (from Sarkar et al 2008, see Methods). Color scale indicate statistical significance of signature overlap (FDR corrected p-values, Fisher's exact test).

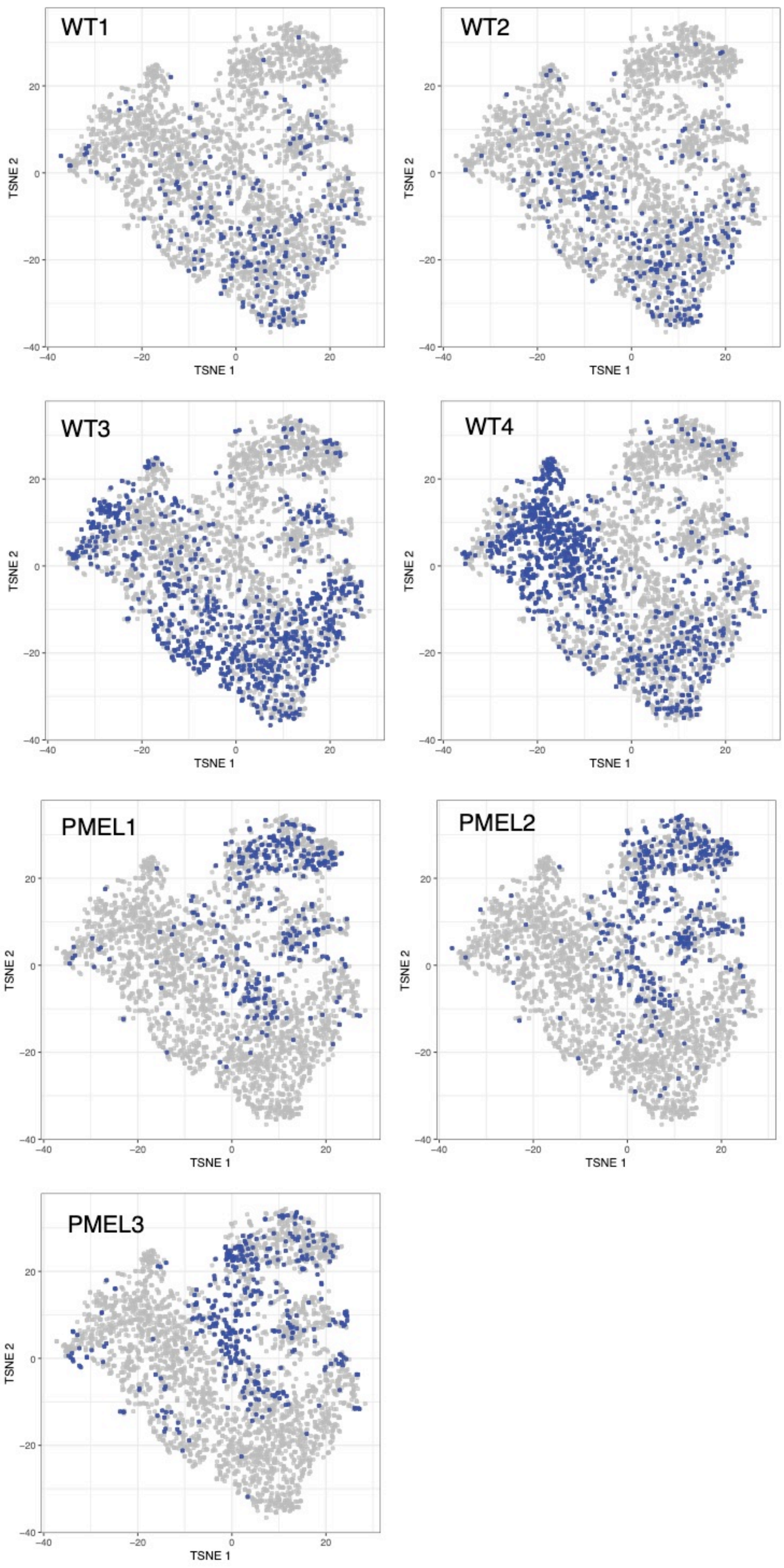


**Supplemental Figure 2** Cell distribution for each individual mouse. Cell from each sample are colored in blue.

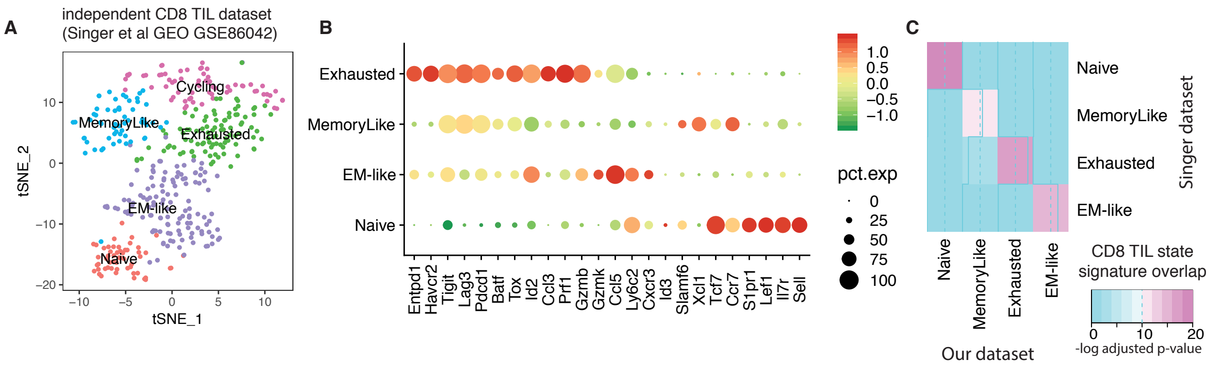


**Supplemental Figure 3** – Unsupervised analysis of an independent CD8 TIL dataset (Singer et al 2016, GEO GSE86042, see Methods). **A** tSNE plot of global transcriptomic similarity of CD8 TILs infiltrating B16 tumors, where colours correspond to the five clusters detected by unsupervised clustering (note that the fifth correspond to a cluster of cycling cells, see Methods). **B** Dotplot indicating average expression of a panel of marker genes (x-axis) for the 4 T-cell subtypes ('cycling' cluster not shown). **C** Heatmap showing cluster similarity (gene signature overlap) between TIL clusters derived from this dataset (Y-axis) and our dataset (C-axis). Color scale indicate statistical significance of signature overlap (FDR corrected p-values, Fisher's exact test).

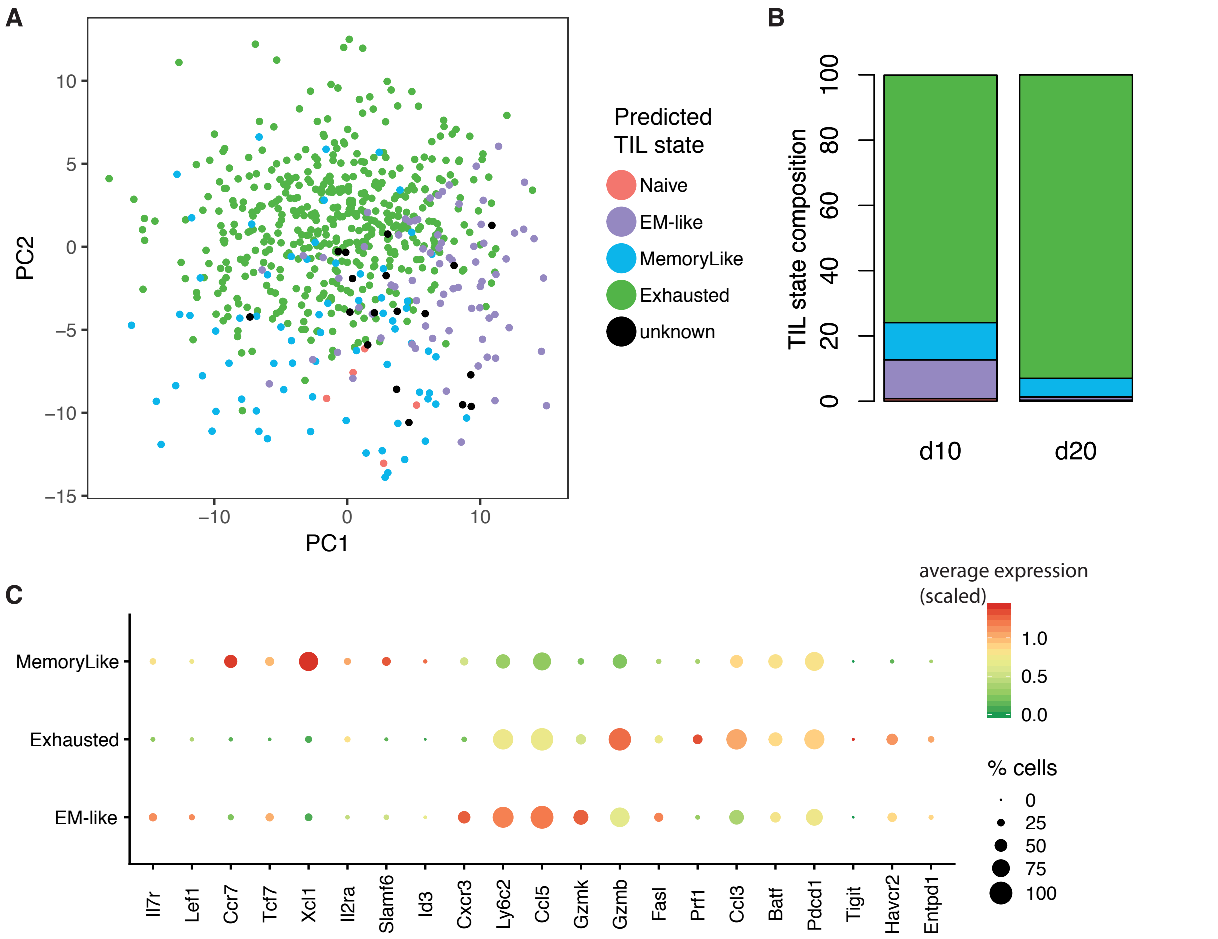


**Supplemental Figure 4** – CD8 TIL state prediction in the (tumor-specific) Tetramer+ CD8 TIL dataset from Miller et al 2019. **A** Tet+ CD8 TILs projected on the first two Principal Components (PCA) of the transcriptomic space (samples from both day 10 and day 20 post tumor engraftment). Points (cells) are colored according to predicted CD8 TIL state (using TILPRED, see Methods). **B** Barplot showing relative proportion of TIL states at day 10 (left) and 20 (right) post tumor engraftment. **C** Dotplot showing average expression of some important markers in the three most abundant predicted states.

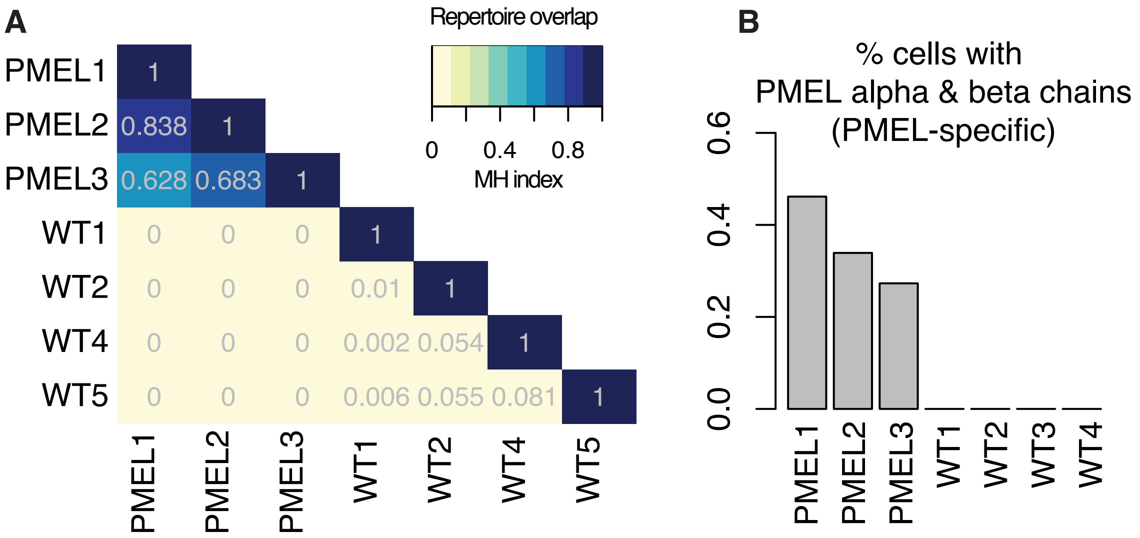


**Supplemental Figure 5** – **A** Clonal overlap between mice (Morisita-Holm index). The large H index between PMEL mice (MH > 0.6) was due to expression of common transgenic PMEL TCR (**B**). The small overlap between wild-type mice was due to the presence of a few public (shared) clones. Further examination revealed that the most abundant public clonotype was TRAV3D-3_TRAJ42_TRAC (CDR3 CAVSEEGSNAKLTF) - TRBV16_TRBD1_TRBJ1−4_TRBC1 (CDR3 CASSHRRANERLFF) shared between mice WT2, WT3 and WT4 (MH 0.05-0.08). This clonotype did not match reported invariant chains and no known epitopes were found for this TCR by literature and database searches. **B** Percentage of cells expressing PMEL TCR (both alpha and beta chains) in each mouse

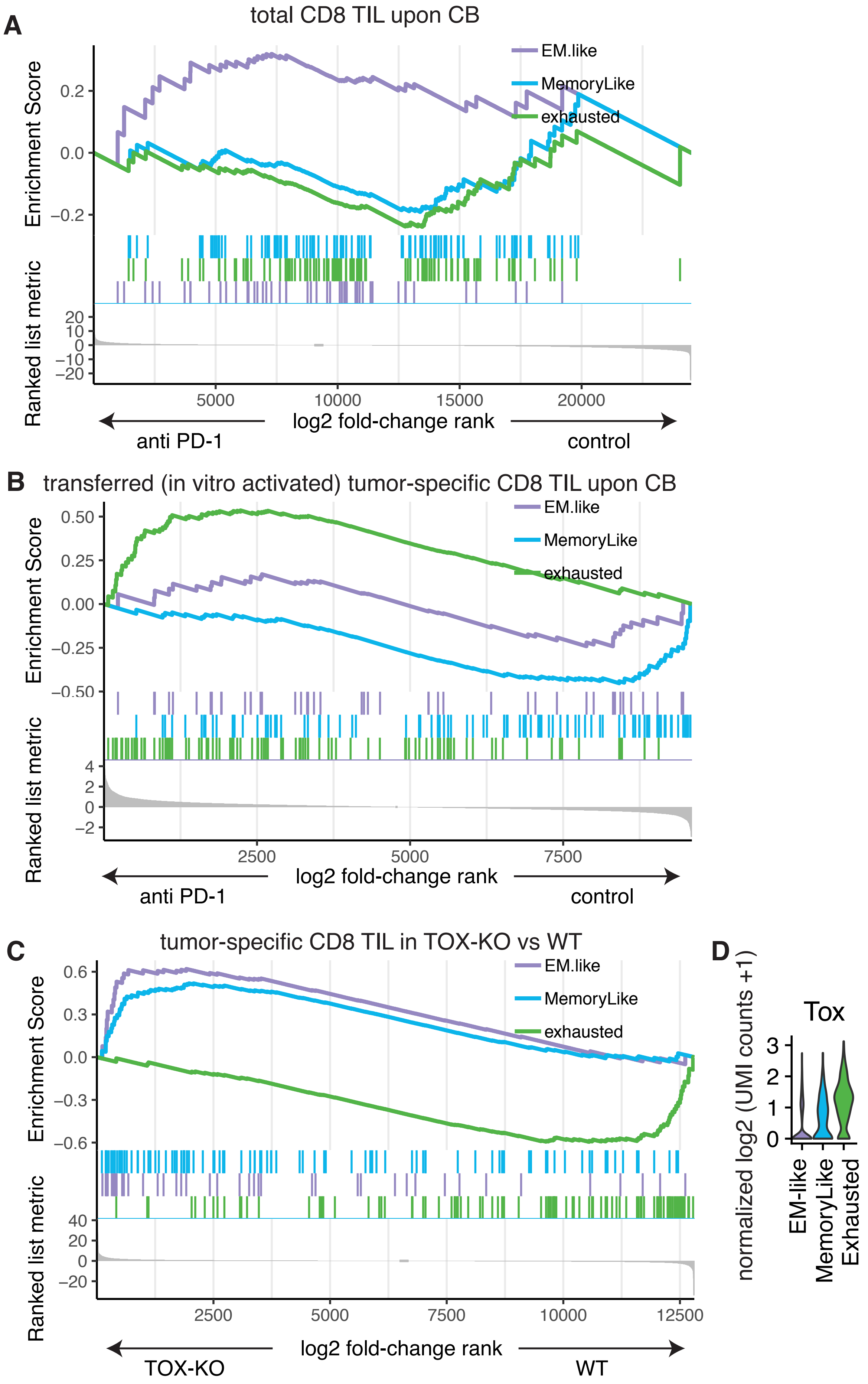
A
total CD8 TIL upon CB
EM.like
MemoryLike
exhausted
Enrichment Score
0.2
0.0
−0.2
Ranked list metric
20
10
0
−10
−20
5000
10000
15000
20000
log2 fold-change rank
anti PD-1
control
B
transferred (in vitro activated) tumor-specific CD8 TIL upon CB
EM.like
MemoryLike
exhausted
Enrichment Score
0.50
0.25
0.00
−0.25
−0.50
Ranked list metric
4
2
0
−2
2500
5000
7500
log2 fold-change rank
anti PD-1
control
C
tumor-specific CD8 TIL in TOX-KO vs WT
EM.like
MemoryLike
exhausted
Enrichment Score
0.6
0.3
0.0
−0.3
−0.6
Ranked list metric
40
20
0
−20
2500
5000
7500
10000
12500
log2 fold-change rank
TOX-KO
WT
D
Tox
normalized log2 (UMI counts +1)
3
2
1
0
EM-like
MemoryLike
Exhausted

**Supplemental Figure 6** – A,B,C Gene-signature enrichment analysis of CD8 T-cells bulk RNA-seq data. **A**. Endogenous CD8 TILs (Non-small cell lung cancer) aPD1 treated vs control mice (Markowitz et al 2018). GSEA Normalized Enrichment Score (NES)= 1.01 (EM-like, p-value>0.1), -0.5 (MemoryLike, p-value>0.1), -0.62 (Exhausted, p-value>0.1). **B**. Adoptively transferred (*in vitro* activated) OT-1 cells infiltrating B16-OVA tumors, aPD-L1 treated vs control mice (Mognol et al 2017). NES= -0.87 (EM-like, p-value>0.1), -1.92 (MemoryLike, p-value=0.004), 1.92 (Exhausted, p-value=0.001). **C** Tumor-specific (SV40 large T cell antigen) CD8 TILs (liver cancer) knockout for TOX (TOX-KO) vs wildtype (Scott et al 2019). NES= 1.69 (EM-like, p-value=0.024), 1.61 (MemoryLike, p-value=0.025), -1.94 (Exhausted, p-value=0.002). **D**. *Tox* expression per TIL state (scRNA-seq data)

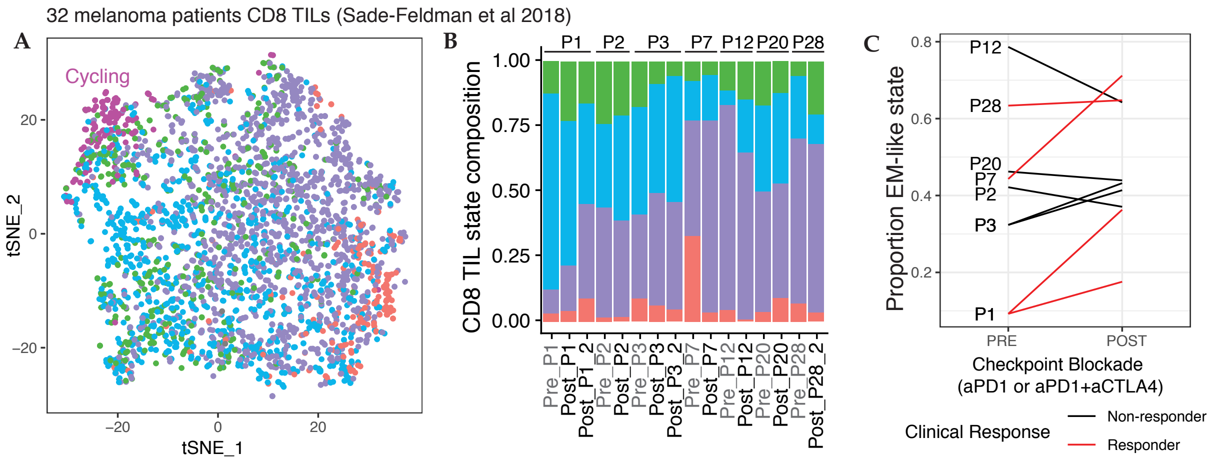


**Supplemental Figure 7** – TILPRED analysis of the CD8 TIL scRNA-seq dataset of Sade-Feldman et al in melanoma patients upon CB

**A** tSNE plot of global transcriptomic similarity of CD8 TILs from 32 melanoma patients (data from (Sade-Feldman *et al.*, 2018)). Cell colors represent TIL state predictions by TILPRED (as in Figure 4 C,D). **B** Proportions of TIL states for selected patients (i.e. patients having samples pre- and post- CB and at least 30 CD8 T cells). Samples are ordered by patients and by timepoint (pre/post CB). **C** Frequency of TILs predicted as EM-like state in samples taken before or after CB. Lines in red for responders and in black for non-responders.
